# Insect egg-killing: a new front on the evolutionary arms-race between Brassicaceae plants and Pierid butterflies

**DOI:** 10.1101/848267

**Authors:** Eddie Griese, Lotte Caarls, Setareh Mohammadin, Niccolò Bassetti, Gabriella Bukovinszkine’Kiss, Floris C. Breman, Erik H. Poelman, Rieta Gols, M. Eric Schranz, Nina E. Fatouros

## Abstract

Evolutionary arms-races between plants and herbivores have been proposed to generate key innovations that can drive diversification of the interacting species. Recent studies reveal that plant traits that target herbivore insect eggs are widespread throughout the plant kingdom. Within the Brassicaceae family, some plants express a hypersensitive response (HR)-like necrosis underneath the eggs of specialist cabbage white butterflies (Pieridae) that leads to eggs desiccating or dropping of the leaf. Here, we studied the evolutionary basis of this trait, its egg-killing effect on and elicitation by specialist butterflies, by screening 31 Brassicaceae species and nine Pieridae species. We show that induction of HR-like necrosis by pierid egg deposition is clade-specific in the economically important Brassiceae tribe (Brassica crops and close-relatives) and in the first-branching genus *Aethionema*. The necrosis is elicited only by pierid butterflies that feed on Brassicaceae plants; four *Pieris* and *Anthocharis cardamines* butterflies, of which the larvae are specialists on Brassicaceae, elicited a HR-like necrosis. Eggs of pierid butterflies that feed on Rhamnaceae (*Gonepteryx rhamni*) or Fabaceae (*Colias* spp.) however, did not elicit such a leaf necrosis. Finally, eggs of *Aglais io*, a species of the sister group Nymphalidae, did not elicit any visible response. Counter-adaptations to HR-like necrosis might have evolved by insect deposition of eggs in clusters or on inflorescences. Our findings suggest that the plants’ egg-killing trait is a new front on the evolutionary arms-race between Brassicaceae and pierid butterflies beyond the well-studied chemical defence traits against caterpillars.

## Introduction

The biodiversity on earth is shaped by numerous factors including inter-organismal interactions that can result in coevolution of adaptive traits. For example, the coevolutionary interactions between plants and insects as described by Ehrlich and Raven^1^ has driven the diversification of plant defensive metabolites^2,3^. In turn, specialist herbivores have evolved detoxification mechanisms, which allow them to feed on their host plants despite these toxic metabolites^4,5^, e.g. monarch butterflies can feed on cardenolide-containing milkweeds^6,7^, and Pieridae and *Plutella* caterpillars on glucosinolate-containing Brassicaceae^8-10^.

The role of plant defences against herbivore eggs has been understudied and underappreciated, especially in a coevolutionary perspective between herbivores and plants. The majority of studies on plant-insect interactions have focused on the feeding life stages of herbivorous insects. Yet, plants can already perceive and respond physiologically to the presence of herbivore eggs before they hatch^11^. The evolution of plant defences against insect eggs is an important first line of defence. In almost half of the ∼400.000 known herbivorous insects, especially in case of lepidopteran and sawfly species, eggs may be the first life stage to come into contact with the targeted host plant. Every insect egg being detected and killed, is one less herbivorous larva or adult insect feeding on the plant in the near future.

Different types of plant defences against insect eggs have been reported in more than thirty plant species including gymnosperms and angiosperms (both monocots and eudicots)^12^. In response to insect egg deposition, plants can produce ovicidal substances^13^, form neoplasms^14,15^ or express a hypersensitive response (HR)-like necrosis beneath the eggs^15-19^. Specifically, HR-like necrosis as an egg-killing defence leading to eggs desiccating and/or falling off the leaf. It has so far been observed in plants of the Pinaceae^20^, Poaceae^21^, Fabaceae^22^, Solanaceae^15,16^ and Brassicaceae^17-19,23^ families. However, the phylogenetic occurrence of the egg-killing trait across these plant families and the phylogenetic co-occurrence in the reciprocal insect pest-clade has yet to be investigated in a similar manner to recent studies of plants and their insect herbivores such as the Brassicaceae plants and Pieridae caterpillars.

Sequence-based phylogenetic analysis^24-26^ has established that the Brassicaceae family is split into a core clade containing 3680 species, sub-divided into three major lineages, and a smaller sister clade containing only the genus *Aethionema* (61 species^27,28^). The model plant *Arabidopsis thaliana* is a representative of Lineage I and the *Brassica* crop plants are representatives of Lineage II. Lineage III is a smaller group mostly restricted to Asia and lacking a model or crop species. Cleomaceae is the sister family of the Brassicaceae^29^. Within the Brassicaceae, defences against feeding herbivores and the genetic basis of this defence have intensively been studied^30-33^. Aliphatic glucosinolates evolved as defensive compounds near or at the origin of the Brassicales clade and became more diverse and complex with plant species radiation. While these compounds play an important role in defending the plants against herbivory, many feeding insects have specialized and evolved effective glucosinolate detoxification and/or excretion mechanisms^8,34-36^.

The Pieridae (whites and sulphurs), containing some 17000 species today, use two major host plants belonging to the Fabales (Fabaceae) and Brassicales (Brassicaceae, Resedaceae, Capparaceae and Cleomaceae); species in some clades also shifted to Rosales (Rhamnacea, Rosaceae) or Santalales^9,37^. Recent phylogenetic reconstruction of the Pieridae indicate that the ancestral host appears to be Fabaceae with multiple independent shifts to other orders. While the Dismorphiinae and nearly all Coliadinae are Fabales feeders, the sister to the Coliadinae, Pierinae, primary feed on Brassicales^38^. The latter thus represent a single origin of glucosinolates feeding^9^. Shortly after the initial evolution of the order Brassicales, some ancestral Pierinae were able to evolve nitrile-specifier proteins (NSPs) that detoxify glucosinolates. This enabled a host shift from their prior Fabaceae hosts to the Brassicales roughly 80 million years ago^9,37^. Similarly, the evolution of glucosinolate sulfatase in *Plutella xylostella* allowed the caterpillar of these moths to feed on Brassicaceae^8^. It has been shown that speciation-rate shifts, as well as genome-duplication events with gene birth-death dynamics, occurred in both Brassicales and Pieridae, usually following a key defence (glucosinolates) or counter-defence (NSPs and sulfatase) invention in one of the coevolutionary partners^37^. To pinpoint the evolution of transitions and innovations, it is necessary to have investigate the trait(s) of interest in a proper phylogenetic context. Defence responses targeting eggs might add a new layer of traits evolved in response to herbivore specialization. Egg-killing responses could then be understood as a first-line-of-defence on top of the later acting glucosinolate defence system.

Eggs of the specialist herbivore *Pieris brassicae* induce HR-like necrosis in the crop plants *Brassica rapa, B. napus* and *Raphanus sativus*^12,39^. However, egg-induced responses have mainly been studied in the black mustard *Brassica nigra* and the model plant *A. thaliana*. On *A. thaliana* egg deposition induces a localized cell death response and higher expression of defence genes resembling HR against pathogens, but a visible necrosis is not expressed and egg-killing never been shown^40,41^. Egg-killing due to a strong necrosis has been shown for the black mustard *B. nigra*. Within *B. nigra*, HR-like necrosis shows high intraspecific variation. Several *B. nigra* accessions were tested with regard to their ability to express HR-like necrosis in response to egg depositions, with some accessions being more likely to express this trait than others^17,18,23^.

The current study explores whether egg-killing necrosis evolved as a specific response to pierid egg deposition in a subset of Brassicaceae. So far, no large-scale screening has been done within the family to determine how common the egg-killing necrosis is expressed within the family. Furthermore, no effort has ever been made to map the phylogenetic history of any egg defence trait for any plant family. Doing so would be a first necessary step to show an adaptive response to egg deposition. For this study we first established that egg wash generated from eggs of *P. brassicae* butterflies and egg deposition on plants yielded a similar plant response on *B. nigra* plants. We then used a representative collection of species in the Brassicaceae (mainly lineage I and II) and three species in the Cleomaceae to investigate the phylogenetic occurrence of egg-killing necrosis across the family. Furthermore, we explored the reciprocal phylogenetic co-occurrence in the Pieridae clade and related species. We compared elicitation of HR-like response by egg deposition and egg wash of three other *Pieris* butterflies (Pierinae) as well as by three relatives, *Anthocharis cardamines* (Pierinae) feeding on *Cardamine* plants of Lineage I, *Colias* spp. (Coliadinae) feeding on Fabaceae and *Gonopteryx rhamni* (Coliadinae) feeding on *Rhamnus* plants belonging to Rhamnaceae. As an outgroup, we used the butterfly *Aglais io* (Lepidoptera: Nymphalidae) that feeds on *Urtica* plants (Urticaceae). We addressed the following questions: (i) Is HR-like necrosis induced in a clade-specific manner within the Brassicaceae? (ii) Is the observed necrosis lowering egg survival under greenhouse and field conditions? (iii) Is elicitation of HR-like necrosis by eggs specific to a particular clade of butterfly species (e.g. genus, subfamily or family) and/or specific to species that co-evolved with the Brassicaceae?

## Material and Methods

### Plants and insects

For our study, we obtained seeds of twenty-eight Brassicaceae and three Cleomaceae species from various sources. The selected plants represent the major lineages in each family. For each plant species, between one and eleven accessions were obtained (Table S1). Per accession, between three and seventeen plants were phenotyped across members of the two families. Two accessions of *B. nigra* (SF48, SF19) were used to assess elicitation of the HR-like necrosis by different butterfly species. Finally, egg-killing was tested for four responsive plant species with the same number of genotypes per species. In preliminary trials, plant species with unknown developmental times were grown to assess their flowering time after sowing. Then, plants were sown in a scheme to ensure similar life stages, i.e. vegetative growth, and sizes if possible. Therefore, plants were between three and six weeks old when being treated with butterfly eggs or egg wash.

For phenotyping the Brassicaceae we used the wash of *Pieris brassicae* eggs. To assess induction of HR-like necrosis on *B. nigra* plants, we used egg deposition from two populations of *P. brassicae, P. napi* L. and *P. rapae* L. and one population of *P. mannii* Mayer (Table S2). Furthermore, we tested egg wash from three populations of *A. cardamines* L., and one population of *G. rhamni* L. and *A. io* L. (Lepidopera: Nymphalidae) (Table S2). Finally, survival was measured for eggs of *P. brassicae, P. napi* and *P. rapae*.

*Pieris brassicae, P. napi* and *P. rapae* were reared on *Brassica oleracea* var. *gemmifera* cv. Cyrus in a greenhouse compartment (21 ± 4°C, 60–80% RH, LD 16: 8). *Pieris mannii* was reared in the same greenhouse, but instead on flowering *Iberis* spp. plants. One population of *A. cardamines* was obtained from a butterfly farm Farma Motyli Zielona Dolina (Babidół, Poland) as hibernating pupae. Hibernation was broken by storing the pupae at 4°C in a cold storage room for five months and another month outdoors. After hibernation, the butterflies were kept in a greenhouse compartment (18± 2°C, 50–60% RH, LD 16: 8) with flowering *Cardamine hirsuta* and *Sisymbrium irio* plants to obtain eggs. *Aglais io* butterflies were kept in cages outside (May to June 2018) with cuttings of *Urtica* sp. plants on which they oviposited. Eggs and/or adults of *A. cardamines, Colias* spp. and *G. rhamni* were also collected outdoors (for locations see table S2); adults were released again when sufficient egg depositions were obtained. *Pieris brassicae* and *A. io* both lay egg clutches, *P. napi* sometimes lays eggs in small groups, while *A. cardamines, G. rhamni, P. mannii* and *P. rapae* lay single eggs.

### Egg wash preparation

Wash from *P. brassicae* eggs was made by fostering females to oviposit on filter paper by pinning the paper to the underside of leaves of *B. oleracea* (Fig. 1a). Within 24 hours after oviposition, the filter paper with the eggs was cut and placed into a 15 ml Falcon tube with purified water (purification system from Millipore Company) at a concentration of 400 eggs per ml. The eggs were left overnight at room temperature. The next morning the supernatant was pipetted off and stored at -20 °C. Before using the egg wash, Tween20 was added at a 0.005 % concentration. The addition of Tween20 was necessary to lower the surface tension of the water droplets, therefore improving the distribution of the egg wash on the waxy leaf surface of some plant species.

**Figure 1:**
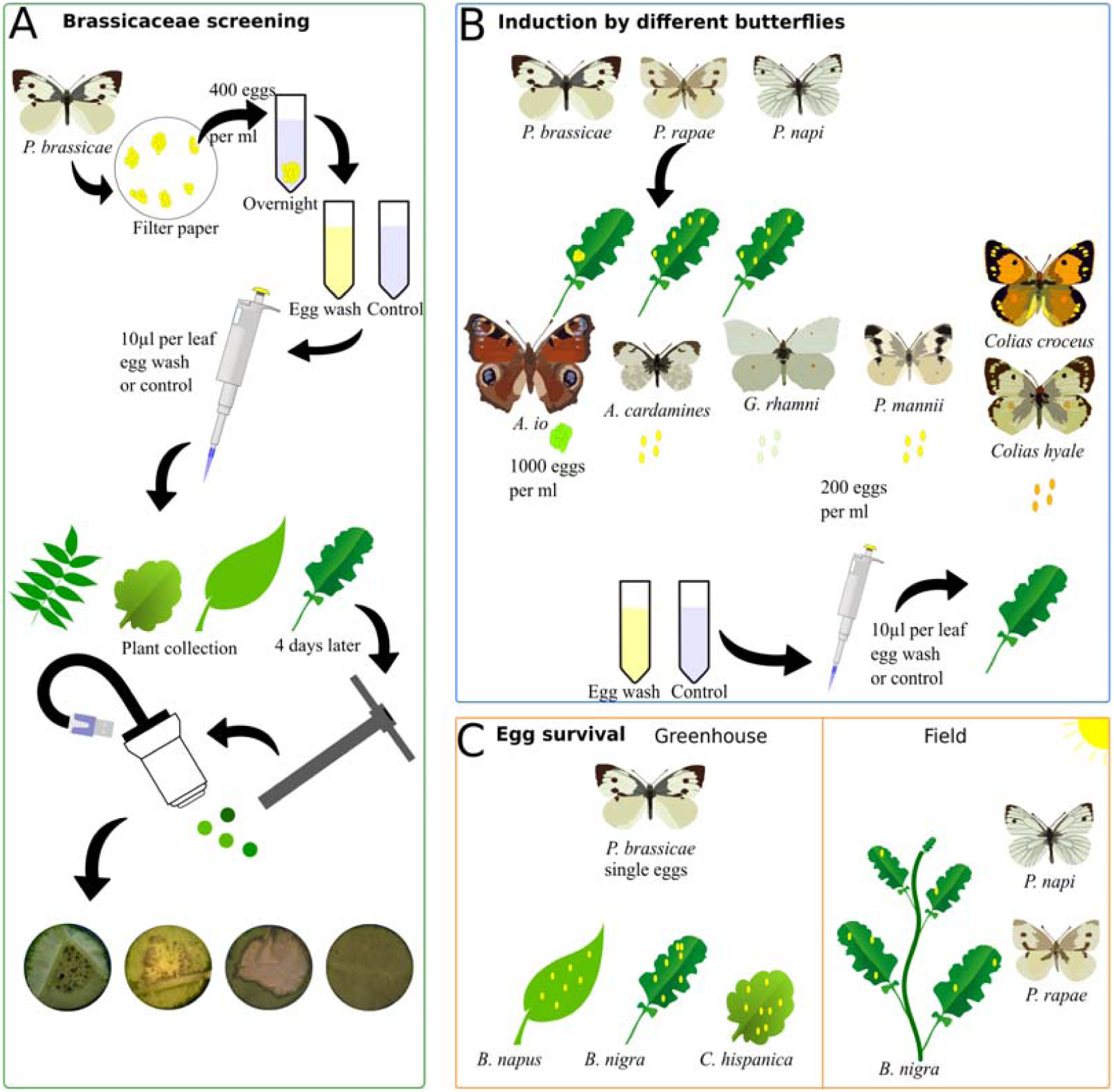
Scheme for plant treatments and phenotyping of HR-like necrosis. A) Production and use of wash from *P. brassicae* egg clusters for a screening of 31 plant species, each of which consisted of 1 to 10 plant accessions. B) Use of eggs or egg wash from different butterfly species to determine which species elicits a necrosis in *B. nigra* accessions. C) Use of singly laid *P. brassicae* eggs to determine the egg-killing effect of HR-like necrosis on *B. napus, B. nigra* and *C. hispanica* accessions. From the field *P. napi* and *P. rapae* eggs were collected from *B. nigra* and hatching (survival) observed.

Wash from *A. io, G. rhamni* and *A. cardamines* eggs was made by removing eggs from leaves of *Urtica* sp. (*A. io*) or *Rhamnus* sp. (*G. rhamni*) and floral inflorescences of *C. hirsuta* or *S. irio* (*A. cardamines*). These eggs were immersed in pure water (*A. io*) or 20 mM 2-(*N*-morpholino) ethane-sulfonic acid (MES) buffer (*A. cardamines*) and left overnight. We chose a concentration of 1000 eggs per ml for *A. io*, as egg size is lower than of Pierini eggs (compare database on egg size from more than 10.000 insect species: https://shchurch.github.io/dataviz/index.html). As controls, clean *Urtica* sp. leaves for *A. io*, a mixture of *C. hirsuta* and *S. irio* inflorescence stems for *A. cardamines*, clean leaves of *Rhamnus frangula* L. for *G. rhamni*, and inflorescence stems of *Iberis* spp. For *P. mannii* were washed in the same manner. Eggs and leaves were kept in the solution overnight, after which the supernatant without eggs was pipetted off and stored at -20 °C. As these egg washes were tested on *B. nigra* plants, no Tween20 was added to the washes.

### Phenotyping of HR-like necrosis of Brassicales plants

Experiments were carried out in a greenhouse compartment to standardize plant-growth conditions (22-27°C, Rh: 50-90%, L:D: 16:8). For the screening of twenty-eight Brassicaceae and three Cleomaceae plant species, 5 µl of *P. brassicae* egg wash was pipetted on a fully mature leaf (the third or fourth leaf from the top) of each plant. Another fully matured leaf (the third or fourth from the top) received pure water with Tween20 as a control. After four days, leaf disks were harvested of the area where egg wash had been applied using a cork borer (1 cm) and put in a rectangular Petri dish with wet blue filter paper. Pictures were taken using a Dino-Lite digital microscope (AnMo Electronics Corporation). These pictures were visually scored for expression of HR-like necrosis (Fig. 1a).

### Testing for elicitation of HR-like necrosis by diverse Pieridae species

Female butterflies of *P. brassicae* (2 populations), *P. napi* and *P. rapae* (2 populations) were allowed to lay between five to ten eggs on two different *B. nigra* accessions (SF19 and SF48) (Supplementary Table 1). Accession SF19 is known as a low responder with respect to egg HR-like necrosis and SF48 as a strong responder^18^. *Anthocharis cardamines, Colias* sp. and *G. rhamni* egg wash was pipetted on both *B. nigra* accessions (Supplementary Table 1). The nymphalid Peacock butterfly *A. io* was used as an outgroup. Eggs laid on *Urtica* leaves were collected and an egg wash made as well as a control wash made from *Urtica* leaves and pipetted on plants of the same *B. nigra* accessions. Between 17 and 40 plant replicates per *B. nigra* accession were used for each butterfly population (Fig. 1b). After four days, HR-like necrosis was scored using a slightly adapted scoring system previously described by Griese et al.^18^. For this scoring system a number between 0 (no response) and 4 (very strong response on both sites of the leaf) is assigned to the observed necrosis.

### Pieris brassicae egg survival on HR-like expressing plants

Experiments were done in greenhouse conditions (21 ± 5°C, Rh: 45 - 70%, L16: D8). HR-like necrosis has been shown to have weaker effects on egg-survival under greenhouse conditions than under natural conditions^17,18^. *Pieris brassicae* females were manipulated to lay five to fifteen separated eggs (not touching each other) on all lines of *B. napus, B. nigra* and *C. hispanica* used in the screening of Brassicaceae species. Previous studies revealed that *P. brassicae* egg survival was only affected when eggs were laid singly, not touching each other^18^. The oviposition of separated eggs was accomplished by observing the females and taking them off the leaf after they laid one egg. After this, the females were put on a different spot of the same leaf. The eggs were left on the plant and four days after oviposition HR-like necrosis was scored as present or absent. After five days, survival of eggs was noted by counting the number of hatched caterpillars (Fig. 1c).

### Pieris brassicae egg survival assessed by field survey

A survey was conducted to record survival of *Pieris* eggs on individual *B. nigra* plants in a natural population (compare Fatouros, et al.^17^). The survey was conducted at an established *B. nigra* patch along the River Rhine in Wageningen (Steenfabriek), The Netherlands (coordinates: 51.96°N, 5.68°E) in one season and butterfly generation (August—September 2017). The total area monitored was approximately 100 m^2^ consisting of ∼1000 plants. Plants were monitored for eggs at the edges of a patch or on isolated growing plants So that not all ∼1000 plants were monitored. Eggs were collected on leaves and checked for the presence of a HR-like necrotic zone on the leaf. After collection, eggs were kept in a climate chamber (25 ± 1°C, 50–70 % RH, L16: D8) until caterpillars emerged. All hatched and dead eggs were recorded (Fig. 1c).

### Phylogenetic analysis of Brassicales and Pieridae species

We used a consensus tree to place our tested Brassicales species according to the species (or genera) reported by two recent studies^25,26^. Both studies analyse representatives of the three distinct linages of the core Brassicaceae clade and the first-branching *Aethionema* and the outgroup Cleomaceae. We used the established three-linage classification when planning and conducting our experiments. As some species and genera were not present in either study, we established their relationships with other included species by calculating our own phylogenetic tree using DNA sequences of two chloroplast markers (*rbcL* and *matK*) and one nuclear genome marker (*ITS2*). The sequences were obtained from the BOLD system website (ID numbers see Supplementary Table 3)^42^. The phylogenetic tree was inferred under maximum likelihood using RaxML v 8.2.4 (GTR+GAMMA, random seed and 1000 bootstrap pseudo-replicates) on the CIPRES science gateway^43,44^. The three Cleomaceae species were used as outgroups for the phylogenetic tree.

The phylogenetic tree of the butterfly species was created using the mitochondrial *COI* gene and the nuclear *EF1*α (Supplementary Table 4). The phylogenetic tree was inferred using maximum likelihood through the IQ TREE website^45-47^. The models selected here for each of the partitions were GTR+F+I+G4:part1, TIM2e+G4:part2, random seed and 1000 ultrafast bootstrap pseudo-replicates. We verified that each clade of butterflies in the tree contained more species than were used in our test to improve separation. *Plutella xylostella* L. was used as an outgroup. The phylogeny showed support for splits within the Pieridae family and the genera were well supported. The phylogeny is very similar to a more extensive study with more species that used two more markers, *wingless* and *28S*^48^.

A Bayesian approach was also performed for phylogenetic inference of the butterflies using the program MrBayes version 3.2^49^ on the same dataset using as priors the parameters from the models selected by IQ TREE and using the same partition of the data. Four simultaneous chains (one cold, three heated) were run for ten million generations, and trees were sampled every 1,000 generations. To check the convergence and stability of the parameter estimates and to determine the burn-in value, Tracer v1.5^50^ was used to explore the log files. Initial trees generated in the burn-in phase (i.e., before establishing stable estimates of parameters) were discarded (burn-in value= 2500, 25 % of the trees). The remaining trees were used to estimate tree topology, branch lengths, and substitution parameters. The phylogenetic relationships inferred from this bayesian approach were congruent with the ML tree obtained from the analysis above.

### Statistical analysis

To test for statistical significance, R version 3.3.2 “Sincere Pumpkin Patch”^51^ was used. For the screening of plant accessions, χ^2^-tests were used to determine which plant species/genotypes significantly expressed HR-like necrosis after egg wash treatment compared to the control treatment. The contingency tables for the χ^2^-tests consisted of the number of egg wash-treated leaves expressing HR-like necrosis, the number of egg wash-treated leaves not expressing HR-like necrosis, the number of control wash-treated leaves expressing HR-like necrosis and the number of control wash-treated leaves not expressing HR-like necrosis. With this set-up, all plant accessions from each plant species were tested independently.

Egg survival was analysed using binomial generalized linear models (GLMs) in which first all variables (plant species, flowering state, HR expression and all interactions between the factors) were used and then based on Akaike information criterions (AICs) removed to simplify the model (plant species, HR expression and interaction). After this, EMMEANS test or Mann-Whitney-U tests were performed as post-hoc tests. Differences in induction of HR-like necrosis by different butterflies were tested using binomial GLMs and, to test differences in strength, GLMs with Poisson distribution Dunn tests with Bonferroni-Holm correction were used as post-hoc tests.

## Results

### Establishing egg wash as an alternative treatment for natural egg deposition

Not all tested butterfly species naturally deposit eggs on (all) brassicaceous species. In order to be able to test eggs of those species and screen a large number of brassicaceous species efficiently, we developed a standard method to wash eggs and treat plants with egg wash. We first compared the effect of eggs and egg wash on *B. nigra*, and scored symptoms induced by oviposition or egg wash, scoring a number between 0 (no response) and 4 (very strong response). The accession SF48 responded with a score between 1-4 in all plants (Supplementary Figure 1). There was no statistical difference between class of symptoms induced by eggs or egg wash (GLM: χ ^2^ = 1.43, df = 1, *P* = 0.232), and so we concluded that we could use egg wash to test the effect on all species.

### Origin of HR-like necrosis in the core Brassicaceae, Aethionema and Cleomaceae

Of all thirty-one species tested, five species responded significantly with HR-like necrosis to *P. brassicae* egg wash. This included species of the genus *Aethionema* and of the tribe Brassiceae (Fig. 2). In the tribe Brassiceae, egg wash treatment significantly enhanced expression of HR-like necrosis in specific accessions of four species: *B. napus* (25-86%), *B. nigra* (63-83%), *B. oleracea* (20-40%) and *C. hispanica* (0-86%) (Supplementary Table 5). There was no significant enhanced HR-like necrosis after egg wash treatment for all other tested plant species tested compared to control leaves. Necrosis was expressed in single plants of some accessions in lineage I and III (0 and 29%) (Fig. 2, Supplementary Table 5). HR-like necrosis of *Aethionema arabicum* varied among the tested accessions between 0 and 60 % (Supplementary Table 5). In some cases, e.g. for *Aethionema carneum*, plants responded with HR-like necrosis to egg wash, however, due to the low number of replicates (*A. carneum*: three plants) difference between control and egg wash treatment was not significant (Supplementary Table 5). For *Lunaria annua*, up to 40% expressed HR-like necrosis, but for this plant species only few replicates were tested, making it impossible to test for significant differences (Supplementary Table 5).

**Figure 2:**
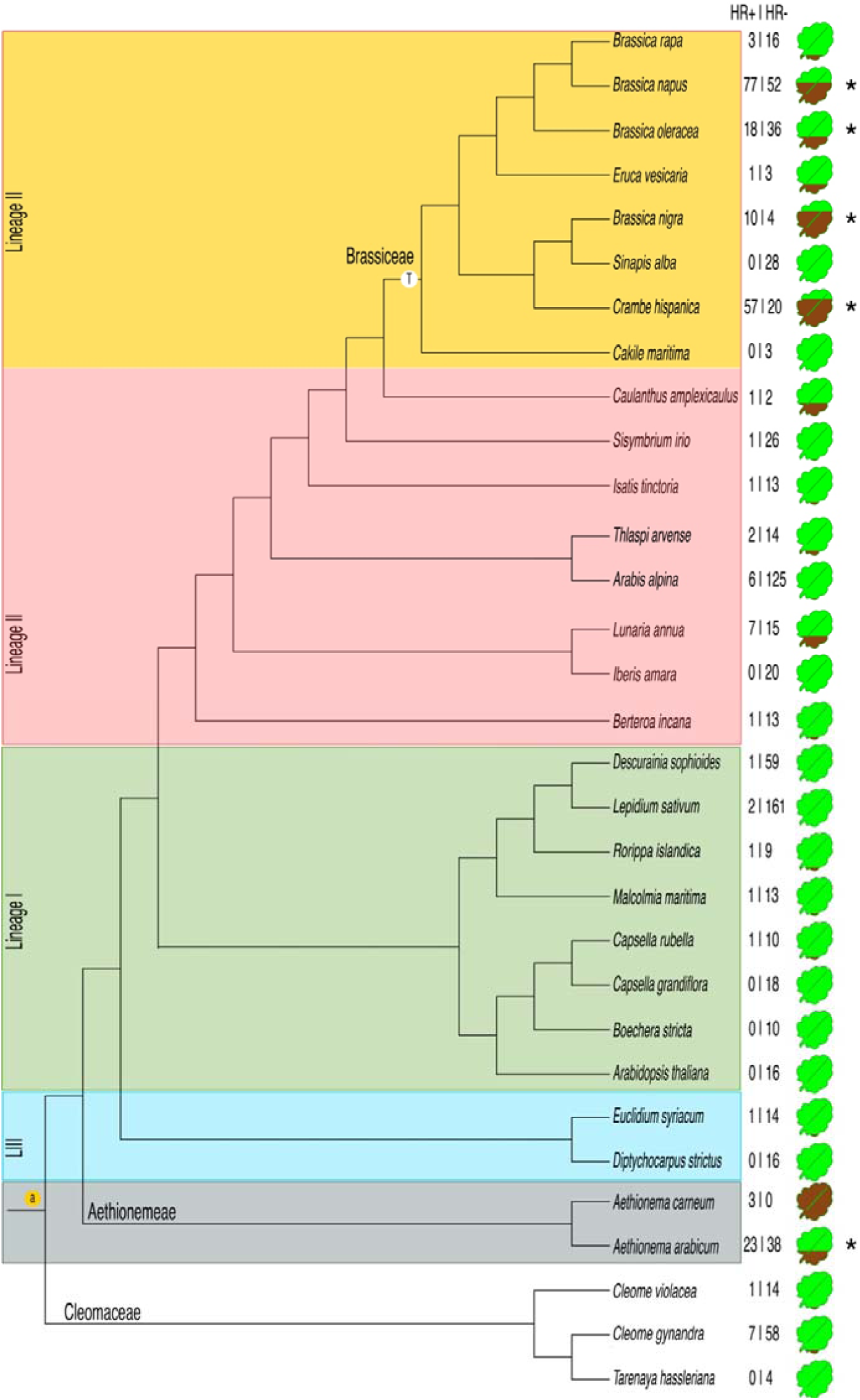
Phylogenetic tree of all plant species treated with *P. brassicae* egg wash and the resulting-fraction of necrosis after 4 days. Consensus phylogeny based on literature and our own analysis of 3 marker genes: rbcL and matK and one nuclear genome marker: ITS2 used. The brown part of the leaf shape represents the percentage of tested plants per plant species responding to egg wash with necrosis. Asterisks indicate that at least one plant accession within the species showed significantly more HR-like necrosis on leaves treated with egg wash than on control treated leaves (χ ^2^-tests, P<0.05). Phylogenetic clades are coloured differently in the tree. The whole genome duplication WGD (a) and genome triplication (T) the Brassiceae tribe specific events are marked in the tree.

### Elicitation of HR-like necrosis by different butterfly species correlated with phylogenetic signal

Egg deposition by all *Pieris* spp. and egg wash of *A. cardamines* elicited a HR-like necrosis on both tested *B. nigra* accessions; the low responding SF19 and as the strong responding SF48. Egg wash of *G. rhamni* and *Colias* spp. did not elicit a HR-like necrosis. Notably, egg wash of both species induced the formation of chlorotic tissue (Fig. 3). Egg wash from *A. io* neither elicited a chlorosis nor HR-like necrosis on either *B. nigra* accession (Table 1). When several populations were available for butterfly species, all populations elicited HR-like necrosis in similar frequency (GLM: χ^2^ = 1.36, df = 3, *P* = 0.71) and severity (GLM: χ^2^ = 2.60, df = 3, *P* = 0.46).

**Table 1:**
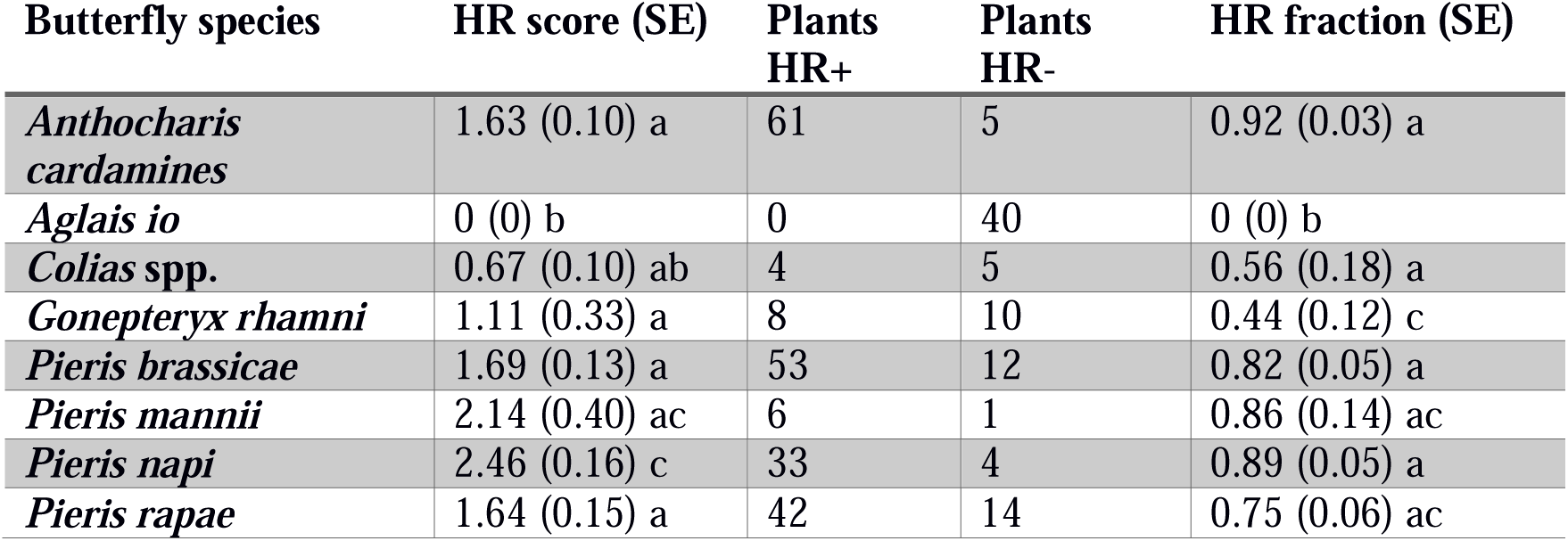
HR-like necrosis (score ranging from 0 to 4) expressed by *B. nigra* plants elicited by different butterfly species. HR- plants did not express HR-like necrosis, while HR+ plants did. Different letters indicate significant differences (different when P < 0.025) between butterfly species, Dunn-test, Bonferroni Holm corrected.

| Butterfly species | HR score (SE) | Plants<br>HR+ | Plants<br>HR- | HR fraction (SE) |
| --- | --- | --- | --- | --- |
| *Anthocharis cardamines* | 1.63 (0.10) a | 61 | 5 | 0.92 (0.03) a |
| *Aglais io* | 0 (0) b | 0 | 40 | 0 (0) b |
| *Colias* spp. | 0.67 (0.10) ab | 4 | 5 | 0.56 (0.18) a |
| *Gonepteryx rhamni* | 1.11 (0.33) a | 8 | 10 | 0.44 (0.12) c |
| *Pieris brassicae* | 1.69 (0.13) a | 53 | 12 | 0.82 (0.05) a |
| *Pieris mannii* | 2.14 (0.40) ac | 6 | 1 | 0.86 (0.14) ac |
| *Pieris napi* | 2.46 (0.16) c | 33 | 4 | 0.89 (0.05) a |
| *Pieris rapae* | 1.64 (0.15) a | 42 | 14 | 0.75 (0.06) ac |

**Figure 3:**
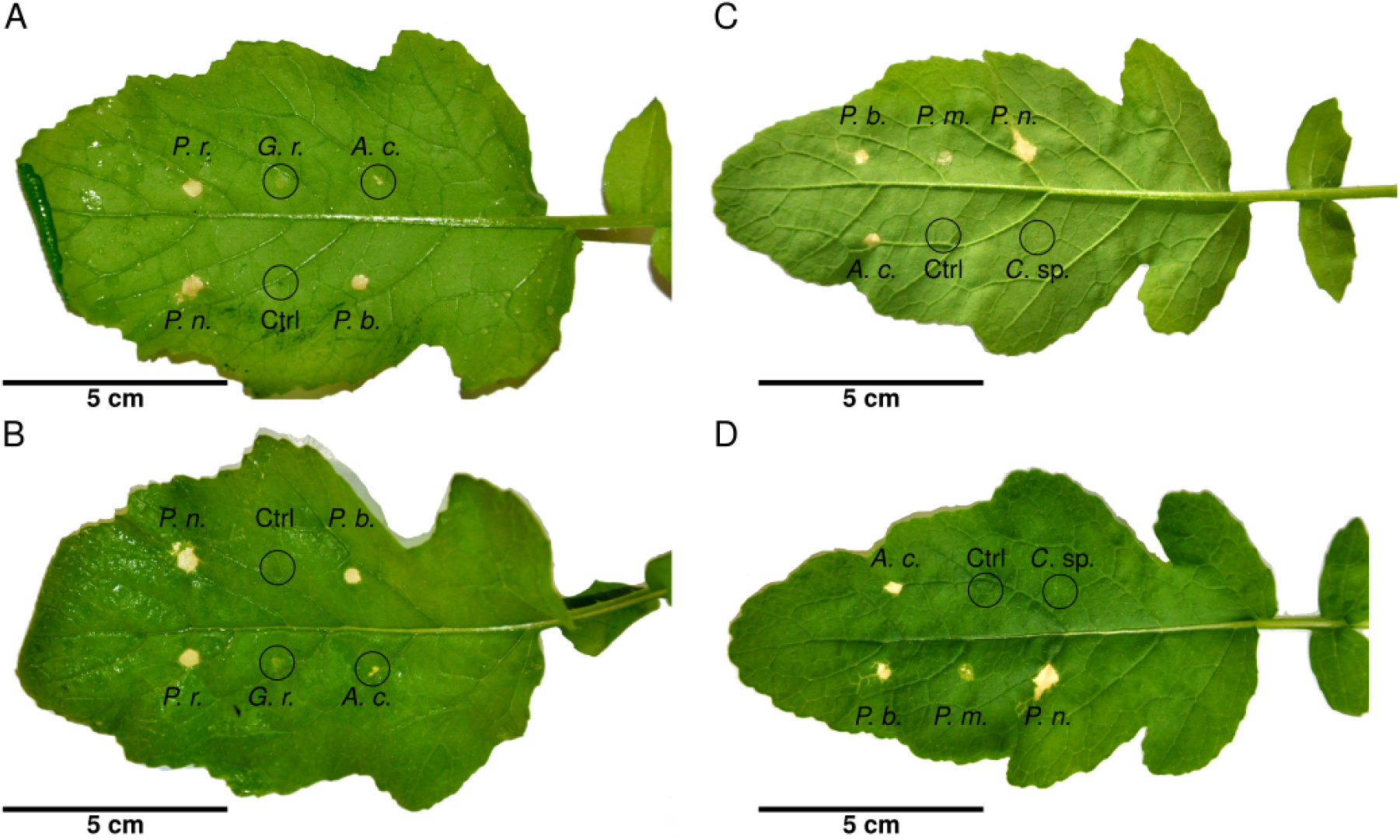
Leaves from *B. nigra* treated with egg wash of different butterfly species and controls inducing or not a HR-like necrosis. *Pieris brassicae* (*P. b.*), *P. mannii*, (*P. m.*), *P. napi* (*P. n.*), and *P. rapae* (*P. r.*) and *Anthocharis cardamines* (*A. c.*) induce a strong HR-like necrosis. Egg wash of *G. rhamni* (*G. r.*) and *Colias* sp. (*C.* sp.) induces a very faint response resembling a chlorosis and does not fit into the established scoring system (faintness indicates 1, but showing up on both sides of the leaf indicates 2). The control (buffer without eggs) does not elicit a HR-like necrosis. All egg washes had the same concentration (200 eggs per ml) and amount applied onto the leaf (5µl). Two leaves were needed as not all egg washes were available at the same time. A) and C) Abaxial side of the leaf where the egg washes were applied onto. B) and D) Adaxial side of the leaf showing how strong the HR-like response is on the side which was not treated with egg wash.

Eggs of all brassicaceous specialists, *Pieris brassicae, P. napi, P. rapae* and *A. cardamines* induced an equally high fraction of HR-like necrosis in *B. nigra* (Supplementary Tables 1 and 6). *Pieris napi* elicited a significantly stronger HR-like necrosis (2.46 ± 0.16) compared to all other butterfly species (Supplementary Tables 1 and 7). The fraction and severity of chlorotic tissue formation elicited by *Colias* spp. and *G. rhamni* was generally lower than HR-like necrosis by the eggs of *Pieris* spp and *A. cardamines* (0.44 ± 0.12; 1.11 ± 0.33 respectively) (Table 1 and Supplementary Tables 6-7). When we plotted the fraction of HR-like necrosis and its severity per butterfly species on our phylogeny, the likelihood and severity of HR-like necrosis is stronger in butterfly species that are the more closely related to *Pieris* sp. (Fig. 4). Thus, all tested Pieridae elicited an egg response while the nymphalid butterfly *A. io* of the sister group never did.

**Figure 4:**
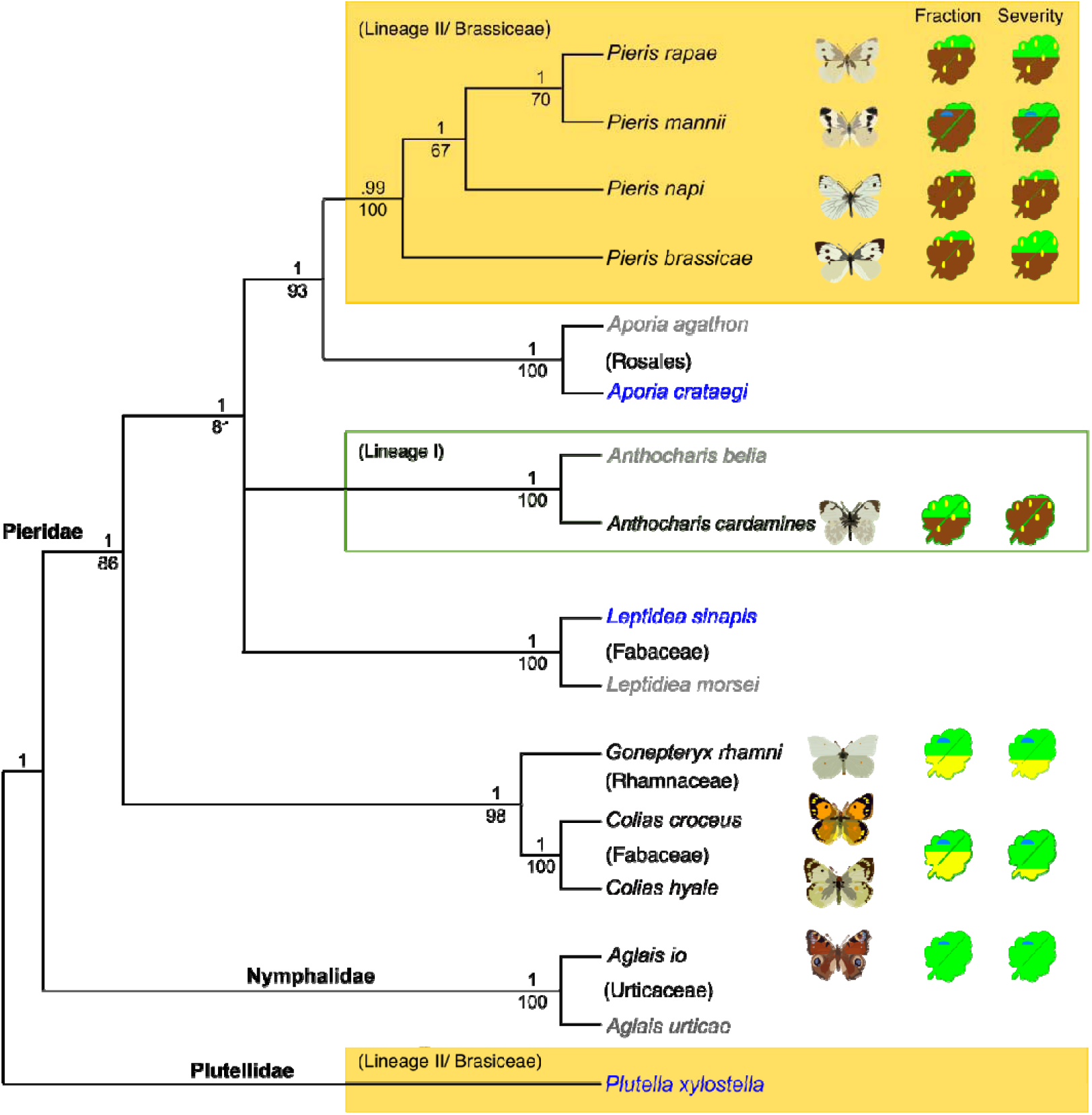
Phylogeny of a subset of Pieridae and elicitation of HR-like necrosis on *B. nigra* leaves by pierid egg wash or eggs. The phylogeny is based on the maximum likelihood and Bayesian posterior probability analysis of the nuclear marker EF1α and mitochondrial maker COI subunit 1. As outgroups, the nymphalid *Aglais io* and the plutellid moth *Plutella xylostella* were chosen. The pictograms of leaves on the right of the cladogram represent the fraction of HR-like necrosis elicitation (left) and severity of HR-like necrosis expressed (right). The average fraction (between 0 and 1) and severity (between 0 and 4) elicited by either eggs or egg wash is represented by the brown part of the leaf, while the yellowing in the leaves represents a different type of response (chlorosis). The phylogenetic tree consists of species used in the experiments (black), species that would answer open questions when tested (blue) and species added to more fully represent the phylogenetic tree (grey). Coloured boxes indicate the Brassicaceae linage which the butterflies use as main host plants. Lepidopteran families are written on their nodes where they separate from the rest of the clades. Bootstrap values for the nodes are given below nodes, Baysian values are given above.

### Effect of HR-like necrosis on Pieris *egg survival on different Brassicaceae plants*

First, we also monitored egg survival of the abundant (in the Netherlands) *Pieris* species (both *P. napi* and *P. rapae*) under natural field conditions. Egg survival was 40 % lower when eggs induced HR-like necrosis compared to survival of eggs that did not induce a leaf necrosis (GLM: χ ^2^ = 11.02, df = 1, P < 0.001, Fig. 5a). As not all eggs on a given plant elicited a necrosis, the fraction of eggs eliciting HR-like necrosis was tested as well.

**Figure 5:**
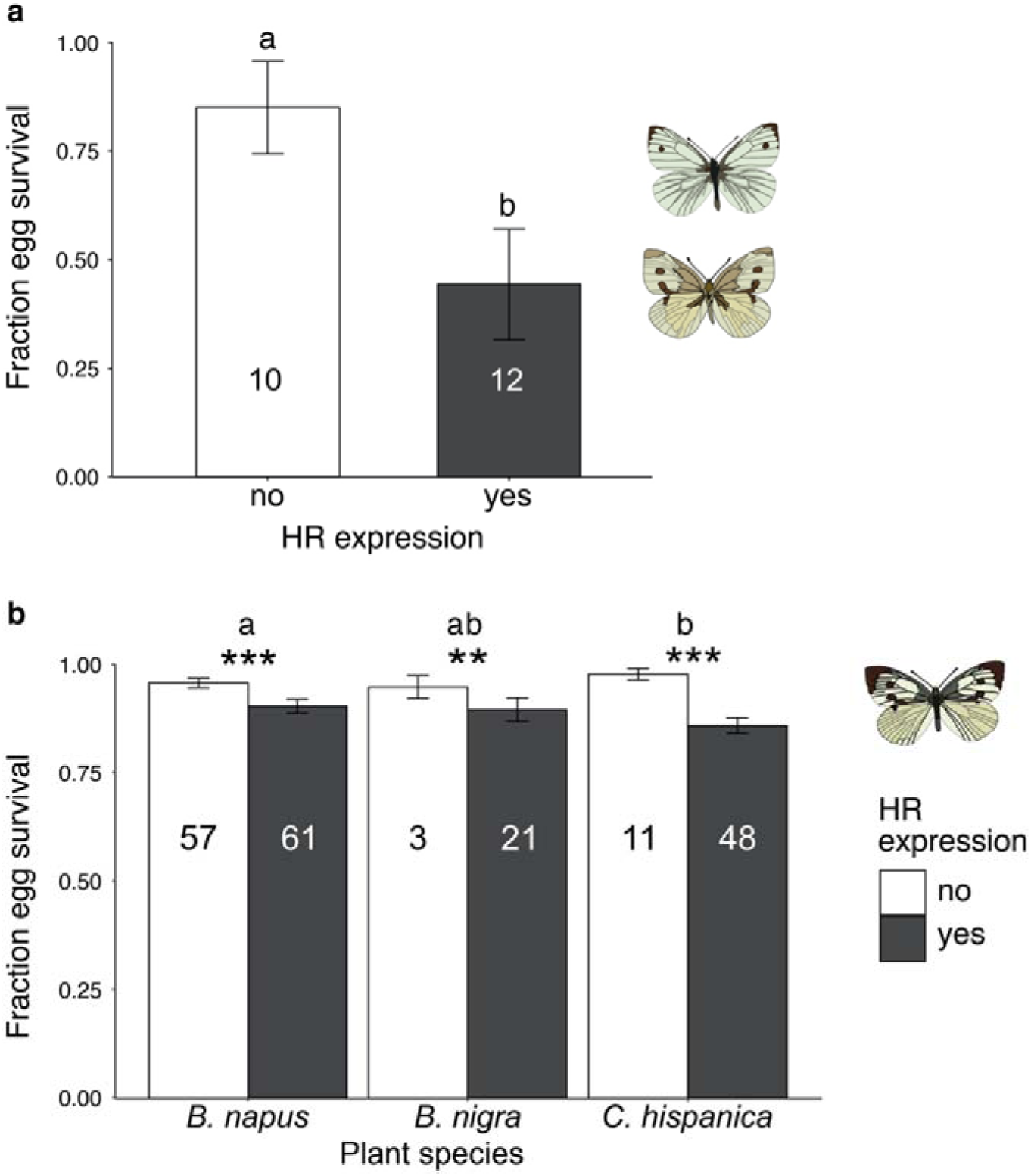
Survival rates of singly laid *P. brassicae, P. rapae* or *P. napi* eggs and effect of expression of HR-like necrosis on different plant species. A) Survey of *P. napi* and *P. rapae* eggs on *B. nigra* plants located near the river Rhine in Wageningen. One to 13 eggs were sampled per plant, total number of collected eggs n = 96. Fraction of survival depending on the expression of HR-like necrosis by the plant. If the plant expressed HR-like necrosis under at least one egg it was counted as HR-expressing ‘yes’. Different letters indicate significant differences (GLM: *P* < 0.001). Numbers in bars indicate the number of plants surveyed within each category. B) Effect of HR-like necrosis on survival rates (mean ± SE) of singly laid *P. brassicae* eggs on *B. napus, B. nigra* and *C. hispanica*. Asterisks indicate differences in egg survival between plants expressing HR-like necrosis and non-HR within a plant species. Different letters indicate significant differences in egg survival between plant species, without taking HR-like necrosis into account. ns: not significant, **: P < 0.01, ***: P < 0.001. (GLM).

Second, we tested egg survival on three highly responding plant species from the first screening under greenhouse conditions. HR-like necrosis significantly lowered the survival of singly laid *P. brassicae* eggs on all three plant species (GLM: χ ^2^ = 38.41, df = 1, *P* < 0.001, fig. 5b). Plant species alone significantly affected egg survival (GLM: χ^2^ = 6.38, df = 2, *P* = 0.04), while the interaction did not (GLM: χ ^2^ = 3.25, df = 2, *P* = 0.20). On *C. hispanica* plants egg survival was significantly lower than on *B. napus* plants (pairwise MWU: *P* = 0.006, Fig. 5b).

## Discussion

Pierid butterflies and their brassicaceous host plants are a fascinating model system of co-evolutionary interactions; research so far has explored its evolutionary and genetic basis by focusing on the diversifying selection on plant chemical defences, i.e. glucosinolates, and insect NSP detoxification genes^9,37,52^. Here, we attempt for the first time to map the phylogenetic history of an egg-induced plant defence trait and its reciprocal co-occurrence in the herbivore clade. We show that pierid egg-induced HR-like necrosis evolved in two clades within the Brassicales. Half of the tested plant species from the Brassiceae tribe in lineage II express strong HR-like necrosis to egg wash. Moreover, all tested *Aethionema* species, the sister clade to the core Brassicaceae, expressed leaf necrosis. Of the *Brassica* and *Crambe* plants (tribe Brassiceae) that were tested, the HR-like necrosis lowered egg survival both under natural and greenhouse conditions. Furthermore, we showed for the first time that only egg wash of *Pieris* butterflies and *A. cardamines*, specialist feeders on the Brassicaceae, elicit a strong HR-like necrosis on *B. nigra*. While *Colias* spp. and *G. rhamni* elicited a chlorotic response similar to that of *Solanum dulcamara* to *Spodoptera* eggs^53^. Our results demonstrate that the egg-induced HR-like necrosis evolved as a new trait at least twice in the Brassicales, but also show that plants specifically evolved this trait to lower egg survival of those pierid species that evolved effective glucosinolate detoxification mechanisms.

Four out of eight tested Brassiceae species, as well as two tested *Aethionema* species showed consistent HR-like necrosis to *Pieris* egg wash in at least one of the genotypes tested. In other plant species, occasionally a single plant showed a light HR-like necrosis. Likely, those plants are false positives, as some plants expressed a light necrosis to control (buffer) wash as well. Alternatively, it could be a general perception response of insect eggs as described for *A. thaliana*^54^. In the latter species it was shown that a lectin receptor kinase, LecRK-I.8, might be involved in early perception of eggs from two widely divergent species, *P. brassicae* and *Spodoptera littoralis*. The ancient genome triplication event in the Brassiceae tribe might have facilitated the evolution of the HR-like necrosis to eggs in this group by increasing the number of resistance genes underlying the trait. Work is underway to identify the genes, which will contribute to a better understanding on the evolution of HR-like necrosis. It is unlikely that the triplication event is the only factor involved in the evolution of HR-like, because *Aethionema* plants respond to *Pieris* eggs with necrosis as well. *Aethionema* species tested here are annuals that occur in dry habitats during a very short time of the year^55^. Interestingly, most tested Brassiceae plants and *Aethionema* are host plants for different *Pieris* butterflies. Both *P. rapae* and *P. napi* eggs are abundant in nature on *B. nigra* and its close relatives like *Sinapis arvensis*^17,19,55,56^. *Pieris ergane* is described to feed on several *Aethionema* species in their south eastern European habitat^57^.

Not all tested plant species within the Brassiceae tribe within Lineage II expressed HR-like necrosis. This could be because we only selected non-responsive genotypes of these plant species or genus. For example, *Sinapis alba*, did not show HR-like necrosis. However, previous work on the close relative *S. arvensis* showed that eggs of *P. rapae* and *P. brassicae* strongly induced HR-like necrosis^39^. This means that that in some genera there is trait variation between species. Alternatively, some plant species might have lost the ability to express HR-like necrosis. Those plants could be less frequently used as host plants for pierid butterflies e.g. because of a phenological mismatch between the plant species and its potential specialist herbivores, as e.g. in the case of *A. thaliana*^58^. In central Europe, *A. thaliana* is usually not attacked by pierid butterflies, as it is rather small and usually completes its life-cycle before caterpillars could develop on the plant^58^. Notably, *A. cardamines* was observed to deposit eggs on *A. thaliana* in North Sweden where both life cycles briefly overlap^59^. Yet, *Pieris* eggs have not been reported to induce a leaf necrosis lowering *Pieris* egg survival on different genotypes of *A. thaliana* including some Swedish accessions^39,40,60^, neither did we observe a visible necrosis on the tested genotype (Col-0) in our experiments when using *P. brassicae* egg wash.

Strong induction of HR-like necrosis seems to be highly specific to *Pieris* butterfly species belonging to the Pierinae clade and feeding on hosts belonging to the Brassiceae clade. Interestingly, another Pierinae species, *A. cardamines*, induced HR-like but feeds on hosts belonging to lineage I of the Brassicaceae (e.g. *Cardamine* sp.^9^). In the latter lineage we did not find species responding with HR-like necrosis. When collecting *A. cardamines* eggs from the inflorescence of *Cardamine* spp. we did not observe any HR-like necrosis (N.E. Fatouros, personal observation). Wash from eggs of species from the non-brassicaceous Coliadinae subfamily, *Colias* spp. and *G. rhamni* and the nymphalid *A. io* did not elicit HR-like necrosis. This suggests that the elicitor for HR-like necrosis is specific for Pierinae butterflies that evolved with Brassicaceae plant species rather than a general molecule present in butterfly eggs. Testing more pierid species from different clades and host plant families is needed to confirm this hypothesis. So far, we also do not know if slight differences of HR-like necrosis elicitation between different *Pieris* species is caused by quantitative differences of the elicitor(s), or by changes in the chemical composition of the elicitor(s). Currently, we are analysing the chemical composition of the egg wash from the different butterfly species to identify the compounds inducing HR-like necrosis.

Previous work has shown that the NSP glucosinolate detoxification gene was a key innovation in the ancestral Pierinae enabling them to shift host plant from Fabaceae to Brassicaceae^9,37^. A recent study revealed another intriguing counter-adaptation to NSP genes: the speciose genus *Erysimum* has recently gained a novel type of chemical defences, the toxic cardenolides. So far, no known specific adaptations to cardenolides have evolved in insect herbivores, including the Pieridae^61^. On the other hand, pierid butterflies may already have found ways to counter-adapt to the egg-killing HR-like necrosis. Clustered eggs of *P. brassicae* were shown to negate the egg-killing effect of the HR-like necrosis^18^. While other advantages of egg clustering have been proposed before^62^, it clearly is helpful in dealing with HR-like necrosis. Although the direct mechanisms of how clustering can protect against egg-killing HR-like necrosis are unknown, it has been shown that desiccation can be slowed down by clustering eggs^18,63^. This might be mitigated by the reduced egg surface area exposed to the environment, compared with single eggs. Other pierid butterflies like *A. cardamines*^64^, *P. mannii* and *P. napi* have been observed to deposit their eggs near or on inflorescence stems of their host plants (N.E. Fatouros, personal observation).

In conclusion, our findings demonstrate that various Brassicaceae plants can mount defences against insect eggs and that these might be under similar selective pressures as plant defences against feeding insects. A coevolutionary arms-race between *Pieris* butterfly eggs and plant species within the Brassiceae clade as well as species within the sister clade *Aethionema* is likely to have occurred. These plants make use of necrotic lesions to lower egg survival and might just have evolved a new mechanism, possibly hijacked from disease resistances, to combat specialist herbivores adapted to their host plants’ toxins. Being a very early, premeditated defence, the mechanism of HR-like necrosis is currently studied as a novel defensive trait to improve resistance of *Brassica* crops against *Pieris* pests.

## Acknowledgements

We thank the employees of Unifarm (Wageningen University and Research) for rearing and caring of the experimental plants used in the experiment. We are thankful to Pieter Rouweler, André Gidding, Frans van Aggelen and Patrick Verbaarschot for rearing the Dutch *Pieris* butterflies used in the experiment. Centre for Genetic Resources, the Netherlands, the Leibniz-Institut für Pflanzengenetik und Kulturpflanzenforschung and BMAP consortium are thanked for the seeds. Furthermore, we thank Prof. Miltos Tsiantis from the Department of Comparative Development and Genetics, Max Planck Institute for Plant Breeding Research for kindly providing *C. hirsuta* seeds, used as host plants for *A. cardamines*. This research has been made possible by funding of the Netherlands Organisation for Scientific Research (NWO) to N.E.F. (NWO/ALW VIDI 14854 and connected Aspasia).

