## Supplementary material for "Insect egg-killing: a new front on the evolutionary arms-race between Brassicaceae plants and Pierid butterflies": Supplemtary data

**Supplementary Table 1**: Used plant species in screening of HR-like necrosis by *P. brassicae* eggs or egg wash. Sources refer to the persons or organisations providing seeds. If studies are cited authors of the study provided seeds and the same plant accessions were used in the referenced study.

| **Family** | **Plant species** | **N genotypes** | **Genotypes** | **Source(s)** |
| --- | --- | --- | --- | --- |
| Brassicaceae | *Aethionema arabicum* | 7 | 84-56-1; 84-56-2; 84-58; Cyp; KE3873;  KE3926OT-01-2; Tur | ^1^ |
|  | *Aethionema carneum* | 1 | KM220087 | ^2^ |
|  | *Arabidopsis thaliana* | 1 | Col-0 |  |
|  | *Arabis alpina* | 9 | ARA; ARAB1; ARAB2; ARAB3; DOR; PAJ; STY; TOT;WCA | ^3^ |
|  | *Berteroa incana* | 1 | Wild | Erik Poelman Laboratory of Entomology, WUR |
|  | *Boechra stricta* | 1 | LTM2 | Michael Eric Schranz, Department of Biosytematics, WUR |
|  | *Brassica napus* | 10 | 4113; 4115; BRA1679; CR185; CR3195; CR3197; CR578; CR671; CR807; CR993 | IPK Gatersleben Genebank |
|  | *Brassica nigra* | 2 | SF19; SF48 | Erik Poelman Laboratory of Entomology, WUR |
|  | *Brassica oleracea* | 4 | K34-3; OH30-3; 14027; W53-4 | ^4^ |
|  | *Brassica rapa* | 3 | BRO-030; RC144; R-o-18 | Guusje Bonnema, WUR Laboratory of Plant Breeding |
|  | *Cakile maritima* | 1 | KM05-0093-10 | BMAP |
|  | *Capsella grandiflora* | 1 | CAPS3 | IPK Gatersleben Genebank |
|  | *Capsella rubella* | 1 | CAPS2 | IPK Gatersleben Genebank |
|  | *Caulanthus amplexicaulus* | 1 | CAB | BMAP |
|  | *Crambe hispanica* | 7 | CR2569; CR2571; CR2572; CR2573; CR2574; CR2578; USDA388853 | IPK Gatersleben Genebank |
|  | *Descurainia sophiades* | 5 | BGH-01; DES1; DES2; DES3; tetraploid | BMAP, IPK Gatersleben Genebank, BMAP |
|  | *Diptychocarpus strictus* | 1 | 05-0397-10-00 | BMAP |
|  | *Eruca vesicaria* | 1 | USDA-PI6333218 | BMAP |
|  | *Euclidium syriacum* | 1 | GC-0587-68 | BMAP |
|  | *Iberis amara* | 2 | IBE1; MK2005-238 | IPK Gatersleben Genebank, BMAP |
|  | *Isatis tinctoria* | 1 |  | Erik Poelman Laboratory of Entomology, WUR |
|  | *Lepidium sativum* | 11 | LEP40; LEP46; LEP51; LEP54; LEP56; LEP57; LEP61; LEP62; LEP63; LEP64; MK2003-169 | IPK Gatersleben Genebank, BMAP |
|  | *Lunaria annua* | 2 | MK2006-705; Wild | BMAP, Erik Poelman Laboratory of Entomology, WUR |
|  | *Malcomia maritima* | 1 | MK2005-111 | BMAP |
|  | *Rorippa islandica* | 1 | GC2006-89 | BMAP |
|  | *Sinapis alba* | 2 | GC0560-79; Wild |  |
|  | *Sisymbrium irio* | 3 | KM88-34-00-20-14; SIS16; SIS4 | BMAP, IPK Gatersleben Genebank |
|  | *Thlaspi arvense* | 1 | GC1211-67 | BMAP |
| Cleomaceae | *Cleoma gynandra* | 7 | gyn100.1; GYN2; INC-06-015; ODS-15-002; TOT-8889; TOT8892; TOT8917 | Biosystematics group WUR, Laboratory of Genetics WUR, Horticulture and Seed Science University of Abomey-Calavi (Benin), German Genebank, World Vegetable Center, Taiwan |
|  | *Cleoma violacea* | 1 | JH-HB813 | BMAP |
|  | *Tarenaya hasslerian* | 1 | 1100 | Michael Eric Schranz, Department of Biosytematics, WUR |

**Supplementary Table 2**: Origin of butterfly populations used in the study.

| **Butterfly species** | **Population** | **Origin** | **GPS** |
| --- | --- | --- | --- |
| *Aglais io* | PL | Butterfly farm in Babidół | 54.2617°N, 18.4418°E |
| *Anthocharis cardamines* | FR | Ban-de-Laveline, Voges | 48.2453°N, 7.0661°E |
|  | FR | Gorges du Segre | 42.4408°N, 2.0803°E |
|  | PL | Butterfly farm in Babidół | 54.2617°N, 18.4418°E |
| *Gonopteryx rhamni* | NL | Wageningen | 51.9864°N, 5.6797°E |
| *Colias sp.* | FR | Montgenevre, Alpes | 44.5533°N, 6.4130°E |
| *Pieris brassicae* | NL | Laboratory of Entomology, Wageningen University | 51.9865°N, 5.6634°E |
|  | PL | Butterfly farm in Babidół | 54.2617°N, 18.4418°E |
| *Pieris napi* | NL | River Rhine, Wageningen | 51.9607°N, 5.6799°E |
|  | FR | Estagel, Pyrenees-Oriental | 42.7724°N, 2.6996°E |
| *Pieris rapae* | NL | River Rhine, Wageningen | 51.9607°N, 5.6799°E |
|  | FR | Estagel, Pyrenees-Oriental | 42.7724°N, 2.6996°E |
| *Pieris mannii* | NL | Wageningen | 51.9705°N, 5.6766°E |

**Supplementary Table 3**: Reference numbers of genes used for generating the phylogenetic tree of Brassicaceae plant species.

|  | **Record number** | | |  |  |
| --- | --- | --- | --- | --- | --- |
| **Species** | **ITS2** | **matK** | **rbcL** | **Database** | **Publication** |
| *Arabidopsis thaliana* | SDH667-14 | GBVE3079-11 | GBVE3091-11 | BOLDSystem |  |
| *Arabis alpina* | MKPCH717-10 | GBVP2280-14 | GBVP2280-14 | BOLDSystem |  |
| *Berteroa incana* | WAT067-12 | GBVT316-13 | WAT067-12 | BOLDSystem |  |
| *Boechera stricta* | BBYUK647-12 | GBVE3157-11 | BBYUK647-12 | BOLDSystem |  |
| *Brassica napus* | SDH677-14 | GBVR4446-13 | GBVR4446-13 | BOLDSystem |  |
| *Brassica nigra* | WAT161-12 | GBVE3189-11 | GBVE3188-11 | BOLDSystem |  |
| *Brassica oleracea* | MKTRT2795-14 | GBVE3190-11 | GBVE3196-11 | BOLDSystem |  |
| *Brassica rapa* | SDH679-14 | GBVE3203-11 | GBVX6649-15 | BOLDSystem |  |
| *Cakile maritima* | ITSAP430-14 | GBVT3310-13 | GBVE3226-11 | BOLDSystem |  |
| *Capsella rubella* | ITSAJ204-14 | GBVE3245-11 | GBVE3244-11 | BOLDSystem |  |
| *Capsella grandiflora* |  | GBVE3236-11 |  | BOLDSystem |  |
| *Caulanthus amplexicaulus* | ITSAP3272-14 |  |  | BOLDSystem |  |
| *Cleome gynandra* | ITSAK4345-14 | GBVE2267-11 | UHURU294-14 | BOLDSystem |  |
| *Cleome violacea* | ITSAK4316-14 | GBVS2993-13 |  | BOLDSystem |  |
| *Crambe hispanica* |  | GBVT3315-13 |  | BOLDSystem |  |
| *Descurainia sophioides* | BBYUK686-12 | FCA174-09 | BBYUK686-12 | BOLDSystem |  |
| *Diptychocarpus strictus* | ITSAJ2329-14 | GBVT283-13 |  | BOLDSystem |  |
| *Eruca vesicaria* | PCCM726-14 | GBVT3325-13 | PCCM726-14 | BOLDSystem |  |
| *Euclidium syriacum* | ITSAF1252-14 |  |  | BOLDSystem |  |
| *Iberis amara* | PCCM703-14 | GBVE3336-11 | GBVE3335-11 | BOLDSystem |  |
| *Isatis tinctoria* | PCCM740-14 | GBVE3341-11 | GBVT4085-13 | BOLDSystem |  |
| *Lepidium sativum* | HIMS329-12 |  | GBVY2605-14 | BOLDSystem |  |
| *Lunaria annua* | MKTRT855-13 | GBVS1366-13 | GBVS934-13 | BOLDSystem |  |
| *Malcolmia maritima* | AM905723.1 |  |  | NCBI | ^5^ |
| *Rorippa islandica* |  | GBVE3455-11 | FCA2965-11 | BOLDSystem |  |
| *Sinapis alba* | SDH747-14 | GBVE3462-11 | GBVE3463-11 | BOLDSystem |  |
| *Sisymbrium irio* | SDH750-14 | GBVE3466-11 | GBVE3467-11 | BOLDSystem |  |
| *Tarenaya hassleriana* | SDH940-14 | GBVP5560-15 | SDH940-14 | BOLDSystem |  |
| *Thlaspi arvense* |  | GBVE3505-11 | GBVU4434-13 | BOLDSystem |  |
| *Aethionema arabicum* | arabicum | arabicum | arabicum |  |  |
| *Aethionema carneum* |  | carneum | carneum |  |  |

**Supplementary Table 4**: Reference numbers of Lepidoptera genes used for generating the Lepidoptera phylogenetic tree.

| **Species** | **Record number** | | **Database** | **Publication** |
| --- | --- | --- | --- | --- |
|  | **COI** | **EF1α** |  |  |
| *Aglais io* | ABOLA874-15 | KY128398.1 | BOLDSystem/NCBI | ^6^ |
| *Aglais urticae* | ABOLD054-16 | AY248811.1 | BOLDSystem/NCBI | ^7^ |
| *Anthocharis belia* | GBGL1322-06 | AY870560.1 | BOLDSystem/NCBI | ^8^ |
| *Anthocharis cardamines* | ABOLD073-16 | LC090568.1 | BOLDSystem/NCBI |  |
| *Aporia agathon* | SHIBU001-13 | AY870590.1 | BOLDSystem/NCBI | ^8^ |
| *Aporia crataegi* | ABOLD067-16 | EU136668.1 | BOLDSystem/NCBI | ^9^ |
| *Colias crocea* | ABOLB043-15 | EF457747.1 | BOLDSystem/NCBI | ^6^ |
| *Colias hyale* | ABOLB009-15 | EF457744.1 | BOLDSystem/NCBI | ^6^ |
| *Gonepteryx rhamni* | EULEP4050-16 | AY870568.1 | BOLDSystem/NCBI | ^8^ |
| *Leptidea morsei* | EZHBA681-07 | GU372654.1 | BOLDSystem/NCBI | ^1^^0^ |
| *Leptidea sinapis* | GBGLP194-13 | AY870573.1 | BOLDSystem/NCBI | ^8^ |
| *Pieris brassicae* | ABOLD049-16 | LC090574.1 | BOLDSystem/NCBI |  |
| *Pieris mannii* | ABOLD642-17 | na | BOLDSystem |  |
| *Pieirs napi* | ABOLD063-16 | AF173401.1 | BOLDSystem/NCBI | ^1^^1^ |
| *Pieris rapae* | ABOLC189-16 | AY870550.1 | BOLDSystem/NCBI | ^8^ |
| *Plutella xylostella* | AGIMP029-14 | KX606845.1 | BOLDSystem/NCBI |  |

**Supplementary Table 5**: Summary of the fraction of necrosis induced by *P. brassicae* egg wash expressed in the plant genotypes tested. The order is alphabetical and does not reflect species relationships. χ2 and P values were generated by Chi-square tests.

| Plant species | Genotype | Fraction necrosis | χ^2^ | *P* |
| --- | --- | --- | --- | --- |
| *Aethionema arabicum* | 84-56-1 | 0.33 | 0.6 | 0.44 |
|  | 84-56-2 | 0.5 | 4.27 | 0.04 |
|  | 84-58 | 0.6 | 10.16 | 0.001 |
|  | Cyp | 0.31 | 2.66 | 0.1 |
|  | KE3873 | 0 | 0 | 1 |
|  | KE3926OT-01 | 0.33 | 0.6 | 0.44 |
|  | Tur | 0.17 | 0 | 1 |
| *Aethionema carneum* |  | 1 | 2.67 | 0.1 |
| *Arabidopsis thaliana* | Col-0 | 0 | 0 | 1 |
| *Arabis alpina* | ARA | 0.11 | 0 | 1 |
|  | ARAB1 | 0.14 | 0.54 | 0.46 |
|  | ARAB2 | 0.07 | 0 | 1 |
|  | ARAB3 | 0.08 | 0 | 1 |
|  | DOR | 0 | 0 | 1 |
|  | PAJ | 0 | 0 | 1 |
|  | STY | 0.1 | 0.002 | 0.96 |
|  | TOT | 0 | 0 | 1 |
|  | WCA | 0 | 0 | 1 |
| *Bertero incana* | -- | 0.07 | 0 | 1 |
| *Boechra stricta* | LTM2 | 0 | 0 | 1 |
| *Brassica napus* | 4113 | 0.86 | 8.14 | 0.004 |
|  | 4115 | 0.43 | 5.3 | 0.02 |
|  | BRA1679 | 0.76 | 17.93 | <0.001 |
|  | CR185 | 0.25 | 1.52 | 0.22 |
|  | CR3195 | 0.86 | 18.54 | <0.001 |
|  | CR3197 | 0.44 | 6.58 | 0.01 |
|  | CR578 | 0.87 | 19.55 | <0.001 |
|  | CR671 | 0.54 | 7.04 | 0.008 |
|  | CR807 | 0.56 | 4.43 | 0.035 |
|  | CR993 | 0.42 | 4.04 | 0.04 |
| *Brassica nigra* | SF19 | 0.63 | 2.4 | 0.12 |
|  | SF48 | 0.83 | 5.49 | 0.02 |
| *Brassica oleracea* | K34-3 | 0.38 | 4.31 | 0.04 |
|  | OH30-3 | 0.375 | 5.13 | 0.02 |
|  | 14027 | 0.4 | 2.81 | 0.09 |
|  | W53-4 | 0.2 | 1.48 | 0.22 |
| *Brassica rapa* | RC144 | 0.17 | 0 | 1 |
|  | RO18 | 0.15 | 0.54 | 0.46 |
| *Cakile maritima* | KM05-0093-10 | 0 | 0 | 1 |
| *Capsella grandiflora* | CAPS3 | 0 | 0 | 1 |
| *Capsella rubella* | CAPS2 | 0.09 | 0 | 1 |
| *Caulanthus amplexicaulus* | CAB | 0.33 | 0 | 1 |
| *Cleome gynandra* | gyn100.1 | 0.25 | 1.52 | 0.22 |
|  | GYN2 | 0.11 | 0 | 1 |
|  | INC-06-01 | 0.5 | 0.67 | 0.41 |
|  | ODS-15-002 | 0 | 0 | 1 |
|  | TOT-8889 | 0 | 0 | 1 |
|  | TOT-8892 | 0.08 | 0.001 | 0.97 |
|  | TOT-8917 | 0 | 0 | 1 |
| *Cleome violaceae* | JH-HB813 | 0.07 | 0.001 | 0.97 |
| *Crambe hispanica* | CR2569 | 0.85 | 15.76 | <0.001 |
|  | CR2571 | 0.86 | 7.29 | 0.007 |
|  | CR2572 | 0 | 0 | 1 |
|  | CR2573 | 0.57 | 8.58 | 0.003 |
|  | CR2574 | 0.87 | 20.46 | <0.001 |
|  | CR2578 | 0.77 | 13.16 | <0.001 |
|  | USDA388853 | 0.69 | 10.88 | <0.001 |
| *Descurainia sophiades* | BGH-01 | 0 | 0 | 1 |
|  | DES1 | 0 | 0 | 1 |
|  | DES2 | 0 | 0 | 1 |
|  | DES3 | 0.06 | 0 | 1 |
| *Descurainia tetraploid* | -- | 0 | 0 | 1 |
| *Diptychocarpus strictus* | 05-0397-10-00 | 0 | 0 | 1 |
| *Eruca vesicaria* | USDA-PI6333218 | 0.25 | 0 | 1 |
| *Eutrema syriacum* | GC-0587-68 | 0.07 | 0 | 1 |
| *Iberis amara* | IBE1 | 0 | 0 | 1 |
|  | IBE11 | 0 | 0 | 1 |
| *Isatis tinctoria* | -- | 0.07 | 0 | 1 |
| *Lepidium sativum* | LEP40 | 0 | 0 | 1 |
|  | LEP46 | 0 | 0 | 1 |
|  | LEP51 | 0.06 | 0 | 1 |
|  | LEP54 | 0 | 0 | 1 |
|  | LEP56 | 0 | 0 | 1 |
|  | LEP57 | 0 | 0 | 1 |
|  | LEP61 | 0.06 | 0 | 1 |
|  | LEP62 | 0 | 0 | 1 |
|  | LEP63 | 0 | 0 | 1 |
|  | LEP64 | 0 | 0 | 1 |
|  | MK2003-169 | 0 | 0 | 1 |
| *Lunaria annua* | Wild | 0.29 | 3.75 | 0.05 |
|  | MK2006-705 | 0.4 | 0.63 | 0.43 |
| *Malcomia maritima* | MK2005-111 | 0.07 | 0 | 1 |
| *Rorippa islandica* | GC2006-89 | 0.1 | 0 | 1 |
| *Sinapis alba* | Wild | 0 | 0 | 1 |
|  | GC0560-79 | 0 | 0 | 1 |
| *Sisymbrium irio* | KM88-34-00-20-14 | 0 | 0 | 1 |
|  | SIS16 | 0.09 | 0 | 1 |
|  | SIS4 | 0 | 0 | 1 |
| *Tarenaya hassleriana* | 1100 | 0 | 0 | 1 |
| *Thlaspi arvense* | GC1211-67 | 0.13 | 0.53 | 0.47 |

**Supplementary Table** **6**: Comparison of the frequency of HR-like necrosis being elicited by different butterfly species on *B. nigra* (Dunn-test with Bonferroni-Holm correction, differences are significant if *P* < 0.025). Z statistic and P-values are given for each comparison of butterfly species.

|  | ***A. io*** | | ***A. cardamines*** | | ***Colias* spp.** | | ***G. rhamni*** | | ***P. brassicae*** | | ***P. mannii*** | | ***P. napi*** | |
| --- | --- | --- | --- | --- | --- | --- | --- | --- | --- | --- | --- | --- | --- | --- |
|  | Z | *P* | Z | *P* | Z | P | Z | *P* | Z | *P* | Z | P | Z | *P* |
| ***A. cardamines*** | 10.03 | < 0.001 |  |  |  |  |  |  |  |  |  |  |  |  |
| ***Colias* spp.** | -3.27 | 0.01 | 2.26 | 0.20 |  |  |  |  |  |  |  |  |  |  |
| ***G. rhamni*** | -3.40 | 0.01 | 3.92 | 0.001 | 0.59 | 1.00 |  |  |  |  |  |  |  |  |
| ***P. brassicae*** | -8.82 | < 0.001 | 1.35 | 0.97 | -1.59 | 0.73 | -3.03 | 0.02 |  |  |  |  |  |  |
| ***P. mannii*** | -4.55 | < 0.001 | 0.37 | 1.00 | -1.30 | 0.97 | -2.01 | 0.33 | -0.23 | 0.82 |  |  |  |  |
| ***P. napi*** | -8.50 | < 0.001 | 0.34 | 1.00 | -1.97 | 0.34 | -3.39 | 0.01 | -0.81 | 1.00 | -0.18 | 0.43 |  |  |
| ***P. rapae*** | -7.88 | < 0.001 | 2.09 | 0.30 | -1.18 | 1.00 | -2.45 | 0.13 | 0.78 | 1.00 | 0.58 | 1.00 | 1.46 | 0.87 |

**Supplementary Table 7**: Comparison of the severity of HR-like necrosis being elicited by different butterfly species on *B. nigra* (Dunn-test with Bonferroni-Holm correction, differences are significant if *P* < 0.025). Z statistic and P-values are given for each comparison of butterfly species.

|  | ***A. io*** | | ***A. cardamines*** | | ***Colias* spp.** | | ***G. rhamni*** | | ***P. brassicae*** | | ***P. mannii*** | | ***P. napi*** | |
| --- | --- | --- | --- | --- | --- | --- | --- | --- | --- | --- | --- | --- | --- | --- |
|  | Z | *P* | Z | *P* | Z | P | Z | *P* | Z | *P* | Z | P | Z | *P* |
| ***A. cardamines*** | 6.83 | < 0.001 |  |  |  |  |  |  |  |  |  |  |  |  |
| ***Colias* spp.** | -1.51 | 0.59 | 2.28 | 0.16 |  |  |  |  |  |  |  |  |  |  |
| ***G. rhamni*** | -3.37 | 0.01 | 1.55 | 0.61 | -0.98 | 0.98 |  |  |  |  |  |  |  |  |
| ***P. brassicae*** | -7.07 | < 0.001 | -0.31 | 1.00 | -2.43 | 0.12 | -1.75 | 0.48 |  |  |  |  |  |  |
| ***P. mannii*** | -4.42 | < 0.001 | -1.11 | 1.00 | -2.49 | 0.11 | -1.92 | 0.36 | -0.98 | 0.82 |  |  |  |  |
| ***P. napi*** | -9.23 | < 0.001 | -3.59 | 0.003 | -4.17 | < 0.001 | -4.00 | < 0.001 | -3.32 | 0.01 | -0.72 | 0.95 |  |  |
| ***P. rapae*** | -6.68 | < 0.001 | -0.08 | 0.47 | -2.30 | 0.16 | -1.57 | 0.63 | 0.21 | 0.83 | 1.07 | 1.00 | 3.41 | 0.01 |

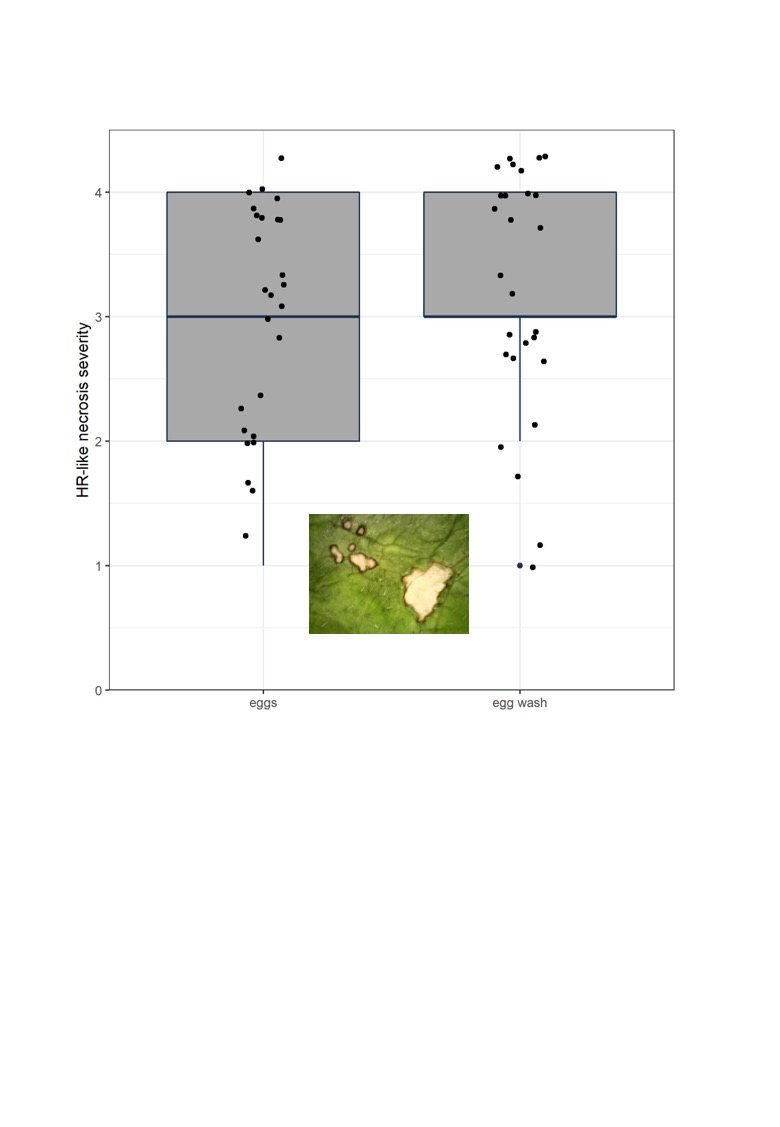

**Supplementary figure 1**: Comparison of the severity of HR-like necrosis induced by *Pieris brassicae* eggs or egg wash in *B. nigra*. Plants were treated with either egg wash or eggs and scored after 72 hours (N=52). There is no statistical difference between the treatments (GLM: χ2 = 1.43, df = 1, P = 0.2315). Inset is a picture of both symptoms induced by eggs (left) and egg wash (right) on the same leaf.
